# A Resource for the Allele-Specific Analysis of DNA Methylation at Multiple Genomically Imprinted Loci in Mice

**DOI:** 10.1101/163832

**Authors:** Jadiel A. Wasson, Onur Birol, David J. Katz

## Abstract

Genomically imprinted loci are expressed mono-allelically dependent upon the parent of origin. Their regulation not only illuminates how chromatin regulates gene expression but also how chromatin can be reprogrammed every generation. Because of their distinct parent of origin regulation, analysis of imprinted loci can be difficult. Single nucleotide polymorphisms (SNPs) are required to accurately assess these elements allele-specifically. However, publicly available SNP databases lack robust verification, making analysis of imprinting difficult. In addition, the allele-specific imprinting assays that have been developed employ different mouse strains, making it difficult to systemically analyze these loci. Here, we have generated a resource that will allow the allele-specific analysis of many significant imprinted loci in a single hybrid strain of *Mus musculus*. This resource includes verification of SNPs present within ten of the most widely used imprinting control regions and allele-specific DNA methylation assays for each gene in a C57BL/6J and CAST/EiJ hybrid strain background.

## Introduction

Genomically imprinted loci, which are expressed mono-allelically dependent upon their parent-of-origin, highlight how DNA methylation and chromatin structure can regulate gene expression (Bartolomei and Ferguson-Smith 2011). For example, many of the chromatin mechanisms that regulate imprinted loci are involved in other contexts, including cancer biology and stem cell reprogramming. In addition, alterations at multiple imprinted loci can be used as a readout of global epigenetic misregulation. As a result, there is an increasing need to assay multiple imprinted loci in different mouse models. In this resource paper, we provide a streamlined resource for assaying the methylation status of a number of the most studied imprinted genes in a single hybrid strain background.

To date, approximately 150 imprinted genes have been identified in mice and about 100 in humans (Gregg *et al.* 2010; DeVeale *et al.* 2012; Kelsey and Bartolomei 2012). These genes tend to be organized on chromosomes in clusters (Wan and Bartolomei 2008; Bartolomei 2009). This clustering allows multiple imprinted loci to be regulated together, under the control of cis-regulatory domains termed imprinting control regions (ICRs) (Wan and Bartolomei 2008; Bartolomei 2009). ICRs are typically between 100 and 3700bp long and are rich in CpG dinucleotides (Bartolomei and Tilghman 1997; Barlow 2011; Ferguson-Smith 2011). In mammals, DNA methylation occurs mainly in the context of CpG dinucleotides, and within ICRs these CpG dinucleotides are differentially methylated dependent upon the parent-of-origin (Reik and Dean 2001; Reik and Walter 2001). This differential methylation determines the expression status of the multiple imprinted genes located within the imprinting cluster (Reik and Walter 2001). Therefore, to globally interrogate the epigenetic control of genomically imprinted loci in a particular mouse model, it is necessary to be able to assay the DNA methylation status of multiple ICRs allele-specifically.

Assessing ICRs allele-specifically requires taking advantage of single nucleotide polymorphisms (SNPs). C57BL/6J (hereafter referred to as B6) mice are the most commonly used strain of *Mus musculus domesticus* and were the first mouse strain to be fully sequenced (Beck *et al.* 2000). To generate hybrids with SNPs on each allele, B6 mice can be crossed to *Mus musculus castaneus* (hereafter referred to as CAST) mice, which originate from a well-defined sub group of wild mice (Beck *et al.* 2000). Genome-wide DNA sequence analysis between different strains of *Mus musculus* revealed a 50% allelic difference between B6 and CAST at potential SNPs (Frazer *et al.* 2007). This makes these hybrid progeny especially useful for analyzing imprinted loci.

SNPs between B6 and CAST are cataloged in the Database of Single Nucleotide Polymorphisms (dbSNP) (http://www.ncbi.nlm.nih.gov/projects/SNP/)(Smigielski *et al.)*2000; Sherry *et al.* 2001). This database reports SNPs that have been observed in various assays performed by individual researchers, consortiums, and genome sequencing centers, for the purpose of facilitating genome-wide association studies (Smigielski *et al.* 2000; Sherry *et al.* 2001). Unfortunately, this database is phasing out all non-human organism data by September of 2017. However, very similar information will still be housed in the European variation archive (http://www.ebi.ac.uk/eva/?Home). This database overlaps with the dbSNP database and also the Sanger SNP viewer database (http://www.sanger.ac.uk/sanger/Mouse_SnpViewer/rel-1505). 2011; Yalcin *et al.* 2011), which provides SNP information in multiple different strain backgrounds.

Using SNP’s from all of these databases, we sought to develop allele-specific DNA methylation assays at multiple ICRs in a B6/CAST hybrid background. However, we encountered two significant hurdles. First, since the dbSNP database and the European variation archive are public repositories, many reported SNPs have not been additionally verified (Mitchell *et al.* 2004; Nekrutenko and Taylor 2012). Moreover, they currently have no minimum requirements for allelic frequencies (Mitchell *et al.* 2004; Nekrutenko and Taylor 2012). This further contributes to the lack of verification for many SNPs. As a result, false positives have been reported at a rate of between 15 and 17 percent (Mitchell *et al.* 2004; Nekrutenko and Taylor 2012). In addition, these two databases pool sequence differences from different strains into one combined output. Thus, we discovered that relying solely on the dbSNP database or European variation archive leads to an even higher rate of false positives within ICRs. These hurdles can partially be overcome by also incorporating the Sanger database, which contains information from individual strain backgrounds. However, a drawback of the Sanger database is that it contains much less information on intergenic regions, where many ICRs are found. For example, it contains no information on 3 of the ICRs that we sought to interrogate. In the end, we assessed 93 B6/CAST SNPs from the three databases at 10 of the most commonly studied mouse ICRs, and were able to validate only 18 of them (19%).

The second hurdle that we encountered is the generation of bisulfite PCR assays within ICRs. The gold standard in probing the DNA methylation status of any locus is bisulfite analysis (Hayatsu *et al.* 2008; Laird 2010). As bisulfite analysis relies on detecting base pair changes at CpG dinucleotides, primer sets used for bisulfite PCR cannot contain any CpG dinucleotides because of the uncertainty of whether a cytosine base in the primer annealing sequence may be methylated. As a result, generating bisulfite-specific primer sets in these highly CpG-rich ICR regions can be difficult. In addition, because the CpG rich ICRs tend to be repetitive, finding primer sets that amplify a unique product can also be challenging.

Based on the significant hurdles we encountered, we identified a need for optimized protocols for allele specific DNA methylation analysis of ICRs in a B6/CAST hybrid mouse background. As a result, we developed a resource, including verification of SNPs present in ICRs, primer information, and optimal PCR conditions. This resource will enable the systematic interrogation of many significant imprinted genes in different mouse models.

## Materials and Methods

### Bisulfite Analysis and Bisulfite-PCR optimization

Mouse Tail DNA from single C57BL/6J and CAST/EiJ animals was used for the original identification of SNPs. Subsequently DNA from sagittal sections of perinatal pups was used for allele-specific DNA methylation analysis. Bisulfite conversion was done according to the Zymo EZ DNA Methylation Kit (Zymo D5001) protocol from 400ng of DNA. PCR products were amplified in a 15μl reaction and 3μl was saved for subsequent TA cloning using the standard TOPO TA cloning protocol (ThermoFisher K4500J10). The remaining volume was run on a 1% agarose gel to confirm that there is a single PCR product. Bisulfite primers were optimized on bisulfite converted DNA using 12 different conditions, including 4 different concentrations of MgCl_2_ (1.5mM, 2.5mM, 3.5mM and 4.5mM) paired with 3 different concentrations of DMSO (0%, 1.5% and 5%). In addition, primers were optimized across a temperature gradient. Primer sets, polymorphisms, and optimal PCR conditions for each gene are listed in the individual figures. Of note, because of the difficulty in finding primer sequences in highly CpG rich regions that do not contain a CpG dinucleotide, many of the primers contained suboptimal base composition and/or did not match the annealing temperature of the other primer used in the reaction. As a result, several of the optimized PCR protocols contain relatively large numbers of cycles to enable the amplification of a product. The BiQ Analyzer program was used for the analysis of bisulfite converted sequences. During the bisulfite analysis, depending on the choice of primers, two different DNA strands will lead to two different sequencing results. Some of the genes we report here were surveyed on the opposite strand of the gene assembly and therefore have a reversed order of their SNPs compared to the databases. These genes are shown with their chromosome location number in reverse order from high to low and this is noted in the corresponding figure legend.

### Data Availability

All data and reagents are available upon request.

## Results

In order to begin the process of interrogating specific imprinted loci, we generated a workflow to streamline the process (Figure 1). Our first criterion was to identify well-defined imprinting control regions (ICRs) that have been extensively studied. We focused on the following ICRs due to their prevalence in the literature: *Grb10, H19, Igf2r, Impact, Lit1/ Kcnq1ot1, Mest/Peg1, Peg3, Peg10, Snrpn*, and *Zac1/Plagl1*. These ICRs also had well-defined locations in the genome and are associated with differentially methylated regions that allowed us to probe their methylation status via bisulfite analysis. We then utilized the UCSC Genome Browser in conjunction with dbSNP to determine reported SNPs within a 10kb window surrounding and including the ICRs, and these SNPs were then crosschecked against the European database, as well as the Sanger database to determine their presence in specific strain backgrounds. Following this *in silico* analysis, we designed bisulfite specific primers to the regions of interest (Table S1). These regions were under 1kb and were within our 10kb defined window, including a significant portion of the ICR and at least one SNP. The bisulfite primers could not contain any CpG dinucleotides, reducing the availability of genomic regions to amplify. Bisulfite primers were optimized on bisulfite converted DNA (detailed in Methods). After optimization, bisulfite PCR was performed on a B6 female and a CAST male, along with the hybrid progeny resulting from the mating. Reported SNPs were compared in B6 and CAST sequences. If validated in this initial comparison, further validation was performed via analysis of the methylation status in hybrid B6/CAST progeny.

Using this workflow, we validated SNPs in all ten ICRs and identified PCR conditions for the analysis of each. The relevant details are reported for each gene below.

### Grb10

*Grb10* is regulated by an ICR that is approximately 1.4kb and located on chromosome 11 in mouse (Figure 2A). Within our probed region, we validated one SNP out of three reported SNPs from the dbSNP database (Figure 2D). The validated SNP is within a 390bp region containing 31 CpG residues (Figure 2A), with the polymorphic base being an A in the B6 background and a G in the Castaneus background (Figure 2B). *Grb10* is methylated on the maternal allele and unmethylated on the paternal allele. This methylation pattern was correctly observed in the hybrid progeny using our optimized assay (Figure 2C and 2E).

**Figure 2:**
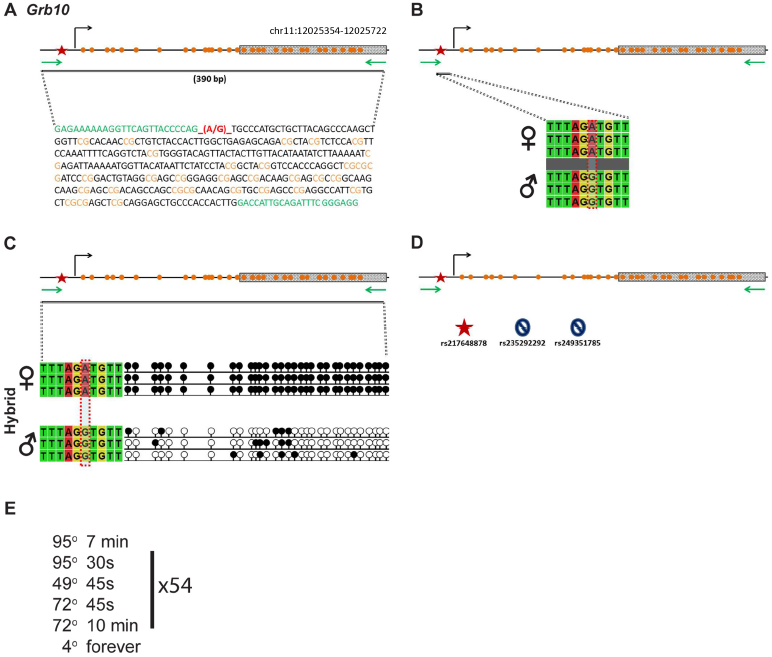
SNP verification within *Grb10* ICR. **A**. Schematic of *Grb10* imprinting control region. Probed region is highlighted by double-dashed line with number of base pairs covered reported, CpG island indicated by dotted box. Green indicates primer sequences orange indicates CpG dinudeotides; red star and bases indicate verified SNP. **B**. Verified SNP presented as sequences from B6 female and CAST male A-to-G SNP is highlighted by red dotted rectangle. **C**. Verification of proper imprinted status in hybrid B6ICAST progeny. SNP highlighted by red dotted rectangle. DNA methylation presented as lollipop diagram; white circles indicate unmethylated cytosines; black cirdes indicate methylated cytosines **D**. Other SNPs reported in all three databases within the probed region with the SNP hlighted by red dotted rectangle. dbSNP identification number indicated under each SNP, Red star indicates validated SNP and blue dosed cirde indicates C-to-T polymorphism that cannot be assayed in bisulfite analysis. **E**. Optimal PCR conditions for probed region with the given primers.

### H19

*H19* is regulated by an ICR on chromosome 7 (Figure 3A). Within our probed region, we validated three SNPs out of four reported SNPs from the dbSNP database (Figure 3D). These validated SNPs are within a 291bp region containing 9 CpG residues (Figure 3A). The three validated SNPs include (1) a G in the B6 background and a deletion in the Castaneus background, (2) a G in the B6 background and an A in the Castaneus background, and (3) an A in the B6 background and a G in the Castaneus background (Figure 3B). *H19* is methylated on the paternal allele and unmethylated on the maternal allele. This methylation pattern was correctly observed in the hybrid progeny using our optimized assay (Figure 3C and 3E).

**Figure 3:**
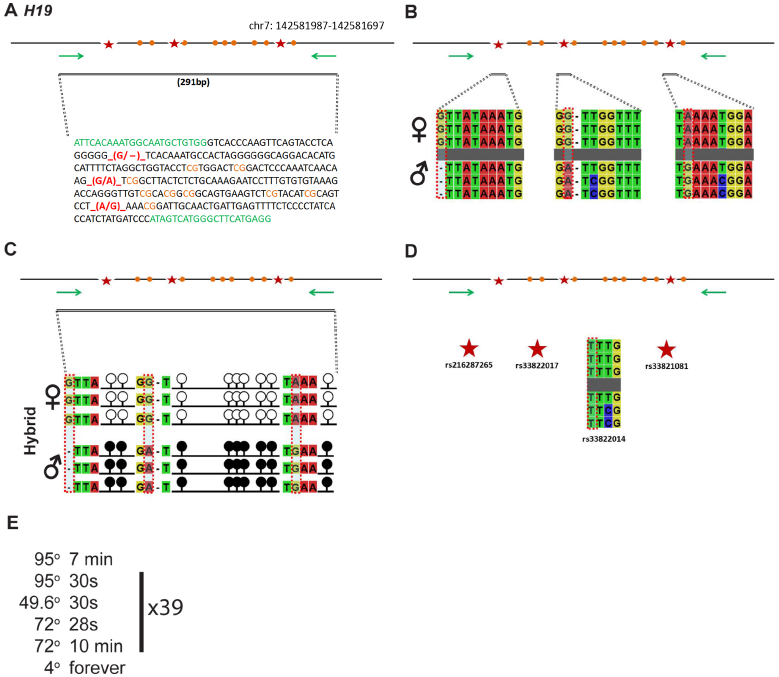
SNP verification within *H19* ICR. **A**. Schematic of *H19* imprinting control region. Probed region is highlighted by double-dashed line with number of base pairs covered reported. CpG island indicated by dotted box. Green indicates primer sequences orange indicates CpG dinucleobdes; red star and bases indicate verified SNPs. The chromosome location is from high to low, see Methds for more details. **B**. Verified SNPs presented as sequences from B6 female and CAST rnaie.G to—del, 0-to-A, and A-to-G SNPs are highlighted by red dotted rectangle. **C**. Verification of proper imprinted status in hybrid 86/CAST progeny. SNPs highlighted by red dotted rectangle. DNA methylation presented as lollipop diagram white circles indicate unmethylated cytosines: black circles Indicate methyLated cytosines. **D**. Other SNPs reported in all three databases within the probed region with the SNP hlighted by red dotted rectangle. dbSNP identification number indicated under each SNP. Red star indicates validated SNP and blue closed circle indicates C-to-T polymorphism that cannot be assayed in bisul. tite analysis. E. Optimal PCR conditions for probed region with the given primers

### Igf2r

*Igf2r* is regulated by an ICR on chromosome 17 (Figure 4A). Within our probed region, we validated two SNPs out of 13 reported SNPs from the dbSNP database (Figure 4D). These validated SNPs are within a 549bp region containing 33 CpG residues (Figure 4A). These polymorphic bases include (1) a G in the B6 background and an A in the Castaneus background, and (2) an A in the B6 background and a G in the Castaneus background (Figure 4B). *Igf2r* is methylated on the maternal allele and unmethylated on the paternal allele. This methylation pattern was correctly observed in the hybrid progeny using our optimized assay (Figure 4C and 4E).

**Figure 4:**
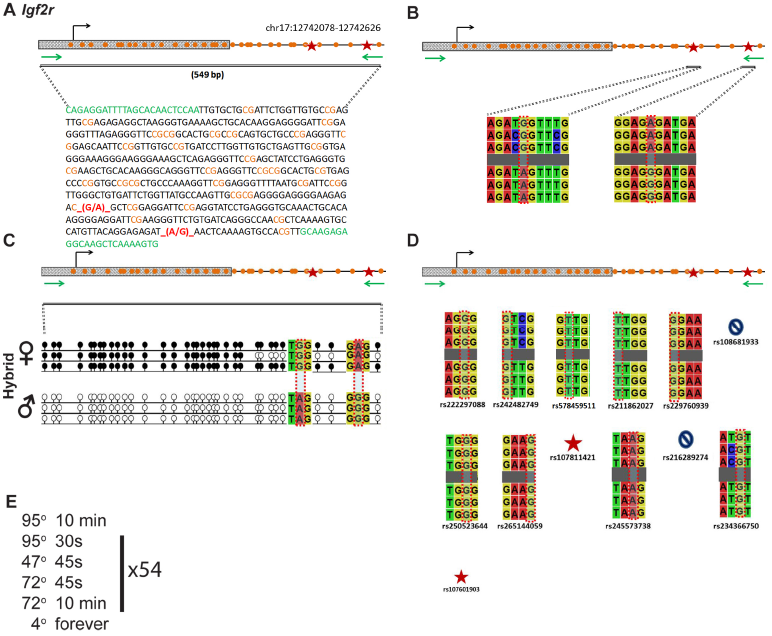
SNP verification within Igf2r ICR. **A**. Schematic of Igf2r imprinting control region. Probed region is highlighted by double-dashed line with number of base pairs covered reported. CpG island indicated by dotted box. Green indicates pnmer sequences; orange indicates CpG dinucleotides; red star and bases indicate venfied SNPs. **B**. Veñfied SNPs presented as sequences from B6 female and CAST male G-to-A and A-to-G SNPs are highlighted by red dotted rectangle, **C**. Verification of proper imprinted status in hybrid B6ICAST progeny. SNPs highlighted by red dotted rectangle. DNA rnethylation presented as lollipop diagram; white circles indicate unmethylated cytosines; black circles indicate methylated cytosines **D**. Other SNPs reported in all three databases within the probed region with the SNP hllghted by red dotted rectangle. dbSNP identification number indicated under each SNP. Red star indicates validated SNP and blue closed circle indicates C-b-T polymorphism that cannot be assayed in bisulfite analysis. **E.** Optimal PCR conditions for probed region with the given primers.

### Impact

*Impact* is regulated by an ICR on chromosome 18 (Figure 5A). Within our probed region, we validated three SNPs out of 10 reported SNPs from the dbSNP and European databases (Figure 5D). One of the SNPs that was not validated was an unnamed SNP from the European database. The validated SNPs are within a 433bp region that contains 17 CpG residues (Figure 5A). These polymorphic bases include (1) a T in the B6 background and an A in the Castaneus background, (2) an A in the B6 background and a G in the Castaneus background, and (3) a T in the B6 background and an A in the Castaneus background (Figure 5B). *Impact* is methylated on the maternal allele and unmethylated on the paternal allele. This methylation pattern was correctly observed in the hybrid progeny using our optimized assay (Figure 5C and 5E).

**Figure 5:**
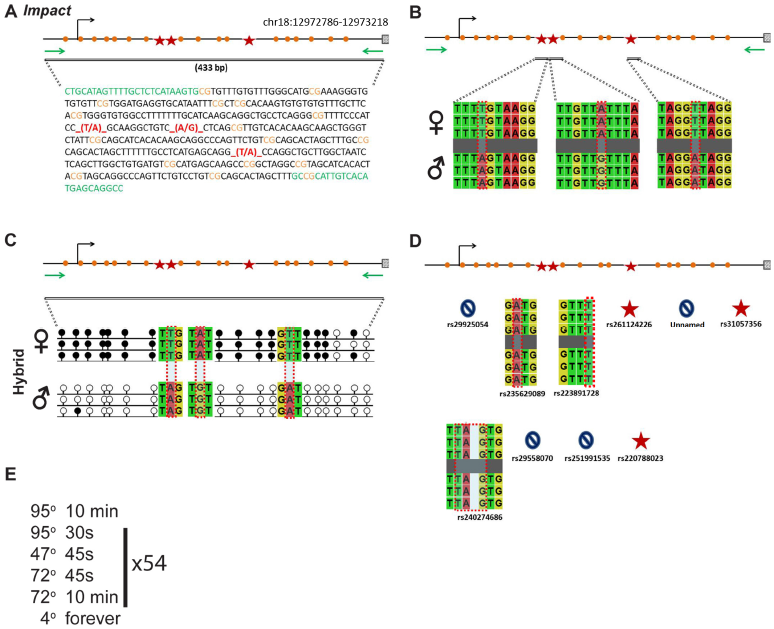
SNP verification within Impact ICR. **A**. Schematic of Impact imprinting contro’ region. Probed region is highlighted by double-dashed line with number of base pairs covered reported. CpG island indicated by dotted box. Green indicates primer sequences; orange indicates CpG dinucleotides; red star and bases indicate verified SNPs. **B**. Verified SNPs presented as sequences from B6 female and CAST male. 110-A, A-to-G. and T-to-A SNPs are highlighted by red dotted rectangle, **C**. Verification of proper imprinted status in hybrid BBICAST progeny. SNPs highlighted by red dotted rectangle. DNA methylation presented as lollipop diagram; white circles indicate unmethylated cytosines: black circles Indicate niethylated cytosines. **D**. Other SNPs reported in all three databases within the probed region with the SNP hlighted by red dotted rectangle. dbSNP identification number indicated under each SNP. Red star indicates validated SNP and blue dosed circle indicates C-to-T polymorphism that cannot be assayed in bisulfite analysis. E. Optimal PCR conditions for probed region with the given primers.

### Lit1/Kcnq1ot1

*Lit1/Kcnq1ot1* is regulated by an ICR on chromosome 7 (Figure 6A). Within our probed region, we validated one SNP out of 12 reported SNPs from the dbSNP and European databases (Figure 6D). One of the SNPs that was not validated was an unnamed SNP from the European database. The validated SNP is within a 420bp region that contains 17 CpG residues (Figure 6A). The polymorphic base is a G in the B6 background and an A in the Castaneus background (Figure 6B). *Lit1* is methylated on the maternal allele and unmethylated on the paternal allele. This methylation pattern was correctly observed in the hybrid progeny using our optimized assay (Figure 6C and 6E).

**Figure 6:**
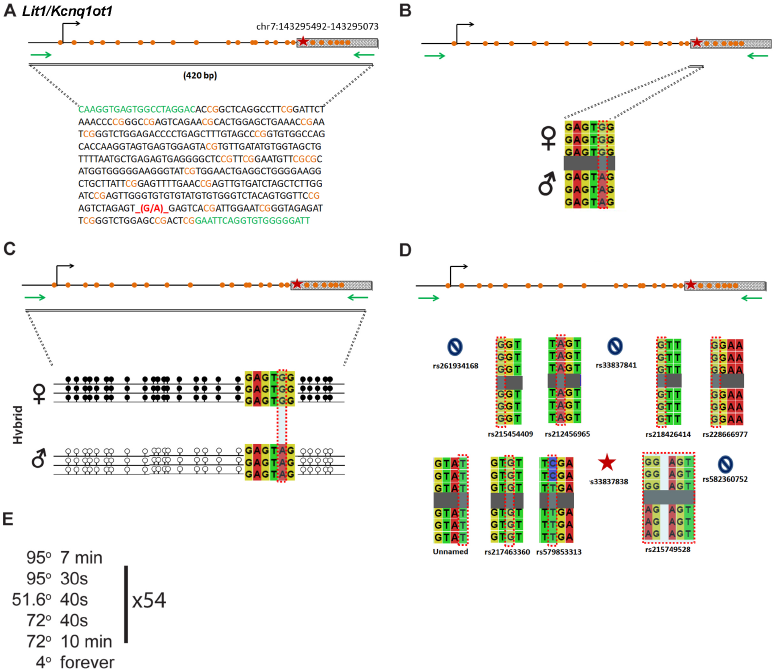
SNP verificatIon within LJtl/Kcnqlotl ICR. **A**. Schematic of Litl/Kcr,q loti imprinting control region. Probed region is highlighted by double-dashed line with number of base pairs covered reported. CpG island indicated by dotted box. Green indicates primer sequences; orange indicates CpG dinucleotides; red star and bases Indicate venhied SNP. The chromosome location Is from high to low, see Methds for more details. **B**. Verified SNP presented as sequences from B6 female and CAST male. G-to-A SNP is highlighted by red dotted rectangle. **C**. Verification of proper imprinted status in hybrid B6/CAST progeny. SNP highlighted by red dotted rectangle. DNA methylation presented as lollipop diagram: white circles indicate unmethylated cytosines; black circles indicate methylated cytosines. **D**. Other SNPs reported in all three databases within the probed region with the SNP hlighted by red dotted rectangle. dbSNP identification number indicated under each SNP. Red star indicates validated SNP and blue closed circle indicates C-to-T polymorphism that cannot be assayed in bisulfite analysis. E. Optimal PCR conditions for probed region with the given primers.

### Mest/Peg1

*Mest/Peg1* is regulated by an ICR on chromosome 6 (Figure 7A). Within our probed region, we validated one SNP out of two reported SNPs from the dbSNP database (Figure7D). This validated SNP is within a 136bp region that contains 4 CpG residues (Figure 7A). This polymorphic base is a T in the B6 background and a G in the Castaneus background (Figure 7B). *Mest* is methylated on the maternal allele and unmethylated on the paternal allele. This methylation pattern was correctly observed in the hybrid progeny using our optimized assay (Figure 7C and 7E).

**Figure 7:**
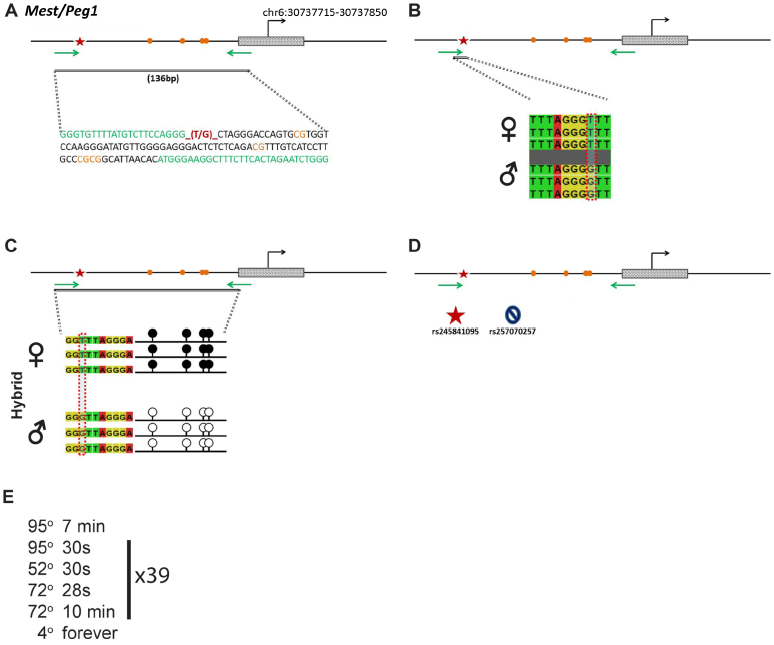
SNP verification within Mest/Pegi ICR. **A**. Schematic of Most/Peg 1 imprinting control region Probed region is highlighted by double-dashed line with number of base pairs covered reported. CpG island Indicated by dotted box Green indicates primer sequences; orange indicates CpG dinucleotides: red star and bases indicate verified SNP. **B**. Verified SNP presented as sequences from B6 female and CAST male T4o-G SNP is highlighted by red dotted rectangle. **C**. Verification of proper imprinted status In hybrid B6/CAST progeny. SNP highlighted by red dotted rectangle. DNA niethylation presented as lollipop diagram: white circles indicate unmethylated cytosines: black circles indicate methylated cytosines. **D**. Other SNPs reported in all three databases within the probed region with the SNP hlighted by red dotted rectangle. dbSNP identification number in dicated under each SNP. Red star indicates validated SNP and blue closed circle indicates C-to-T polymorphism that cannot be assayed In bisulfite analysis. **E**. Optimal PCR conditions for probed region with the given primers.

### Peg3

*Peg3* is regulated by an ICR on chromosome 7 (Figure 8A). Within our probed region, we validated one SNP out of four reported SNPs from the dbSNP database (Figure8D). This validated SNP is within a 228bp region that contains 11 CpG residues (Figure 8A). This polymorphic base is a T in the B6 background and a G in the Castaneus background (Figure 8B). *Peg3* is methylated on the maternal allele and unmethylated on the paternal allele. This methylation pattern was correctly observed in the hybrid progeny using our optimized assay (Figure 8C and 8E).

**Figure 8:**
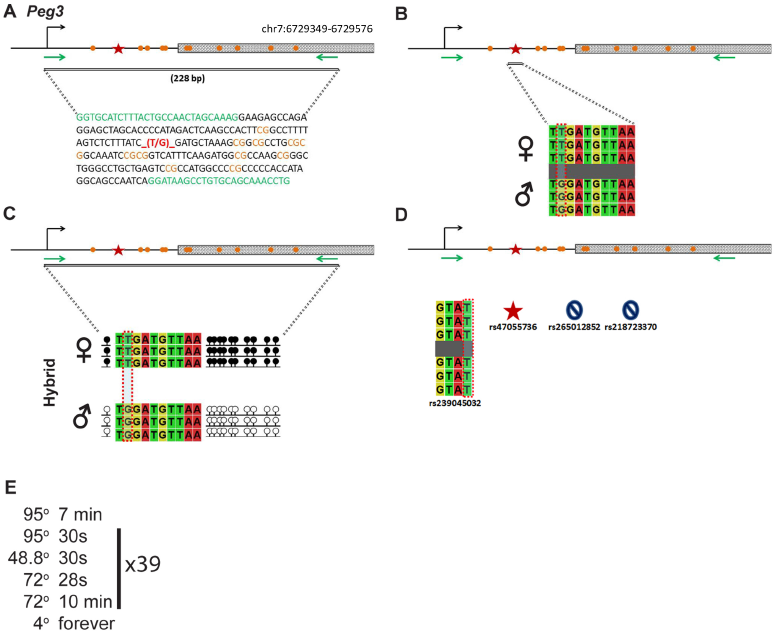
SNP verification within Peg3 ICR. **A**. Schematic of Peg3 imprinting control region. Probed region is highlighted by double-dashed line with number of base pairs covered reported, CpG island indicated by dotted box Green indicates primer sequences; orange indicates CpG dinucleotides; red star and bases indicate verified SNP. B. Verified SNP presented as sequences from B6 female and CAST male T-to-G SNP is highlighted by red dotted rectangle. **C**. Verification of prnper imprinted status in hybrid B6ICAST progeny. SNP highlighted by red dotted rectangle, DNA methylation presented as lollipop diagram; white circles indicate unmethylated cytosines; black circles indicate methylated cytosines. **D**. Other SNPs reported in all three databases within the probed region with the SNP hllghted by red dotted rectangle. dbSNP Identification number indicated under each SNP, Red star indicates validated SNP and blue closed circle indicates C-to-T polymorphism that cannot be assayed in bisulfite analysis, **E**. Optimal PCR conditions for probed region with the given primers.

### Peg10

*Peg10* is regulated by an ICR on chromosome 6 (Figure 9A). Within our probed region, we validated one SNP out of 23 reported SNPs from the dbSNP and European databases (Figure 9D). One of the SNPs that was not validated was an unnamed SNP from the European database. The validated SNP is within a 663bp region that contains 54 CpG residues (Figure 9A). This polymorphic base is a C in the B6 background and an A in the Castaneus background (Figure 9B). *Peg10* is methylated on the maternal allele and unmethylated on the paternal allele. This methylation pattern was correctly observed in the hybrid progeny using our optimized assay (Figure 9C and 9E).

**Figure 9:**
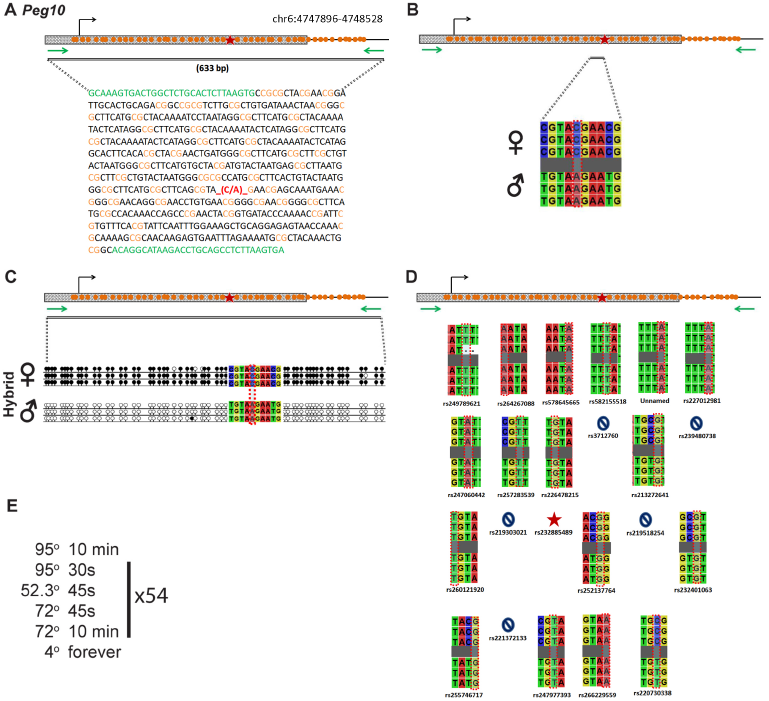
SNP verification within PeglO ICR. **A**. Schematic of PeglO imprinting control region. Frobed regron is dashed line with number of base pairs covered reported. CpG island indicated by dotted box. Green indicates primer sequences; orange indicates CpG dinucleotides; red star and bases indicate verified SNP. **B**. Verified SNP presented as sequences from B6 female and CAST male. C-to-A SNP is highlighted by red dotted rectangle. **C**. Verification of proper imprinted status in hybrid B6/CAST progeny. SNP highlighted by red dotted rectangle. DNA methylation pi-esented as lollipop diagram; white circles indicate unmettiylated cytosines; black circles indicate methylated cytosines. **D**. Other SNPs reported in all three databases within the probed region with the SNP hlighted by red dotted rectangle. dbSNP identification number indicated under each SNP. Red star indicates validated SNP and blue closed circle indicates C-to-T polymorphism that cannot be assayed in bisulfite analy sis. E. Optimal PCR conditions for probed region with the given primers.

### Snrpn

*Snrpn* is regulated by an ICR on chromosome 7 (Figure 10A). Within our probed region, we validated four SNPs out of 11 reported SNPs from the dbSNP database (Figure10D). We also identified a novel SNP that is not present in any of the three databases. All five of the validated SNPs are within a 356bp region that contains 16 CpG residues (Figure 10A). These polymorphic bases include (1) a T in the B6 background and an G in the Castaneus background. This is the novel SNP that we identified. (2) a TTT in the B6 background and a deletion in the Castaneus background, (3) a T in the B6 background and an A in the Castaneus background, (4) a G in the B6 background and an A in the Castaneus background, and (5) a G in the B6 background and a T in the Castaneus background (Figure 10B). *Snrpn* is methylated on the maternal allele and unmethylated on the paternal allele. This methylation pattern was correctly observed in the hybrid progeny using our optimized assay (Figure 10C and 10E).

**Figure 10:**
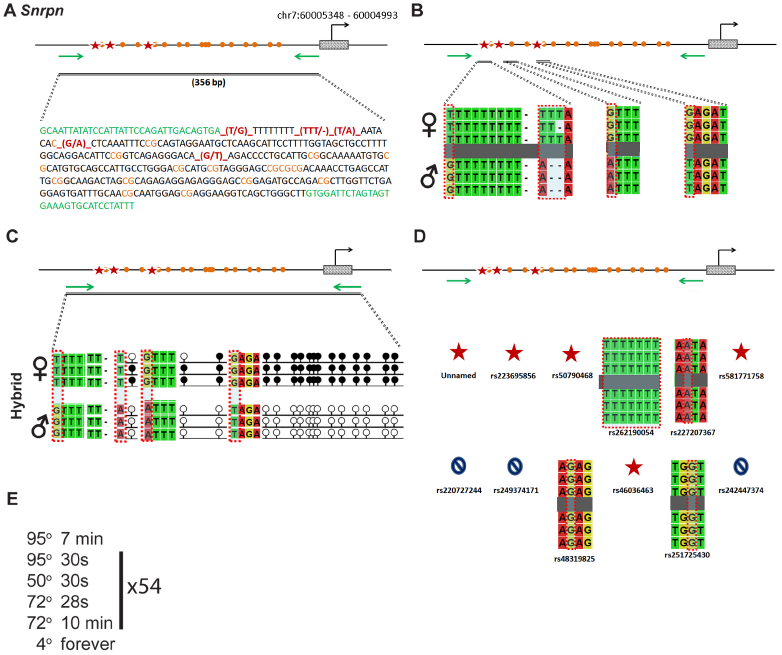
SNP verification within Snrpn ICR. **A**. Schematic of Snrpn imprinting control region. Probed region is highlighted by double-dashed line with number of base pairs covered reported. CpG island indicated by dotted box. Green Indicates primer sequences; orange indicates CpG dinucleotides: red star and bases indicate verified SNPs. The chromosome location is from high to low, see Methds for more details. **B**. Verified SNPs presented as sequences from B6 female and CAST male. T-to-G, TU-to-Del. T-to-A, G-to-A, and G-to-T SNPs are highlighted by red dotted rectangle. **C**. Verification of proper imprinted status In hybrid B6ICAST progeny. SNP highlighted by red dotted rectangle. DNA methylation presented as lollipop diagram: white circles indicate unmethylated cytosines: black circles indicate methylated cytosines. **D**. Other SNPs reported in all three databases within the probed region with the SNP hlighted by red dotted rectangle. dbSNP identification number indicated under each SNP. Red star indicates validated SNP and blue closed circle indicates C-b-T polymorphism that cannot be assayed in bisulfite analysis. E. Optimal PCR conditions for probed region with the given primers.

### Zac1/Plagl1

*Zac1/Plagl1* is regulated by an ICR on chromosome 10 (Figure 11A). Within our probed region, we validated one SNP out of 11 reported SNPs from the dbSNP and European databases (Figure11D). The unnamed SNPs are not found in the dbSNP. The validated SNP is within a 578bp region that contains 33 CpG residues (Figure 11A). This polymorphic base is an A in the B6 background and a G in the Castaneus background (Figure 11B). *Zac1* is methylated on the maternal allele and unmethylated on the paternal allele. This methylation pattern was correctly observed in the hybrid progeny using our optimized assay (Figure 11C and 11E).

**Figure 11:**
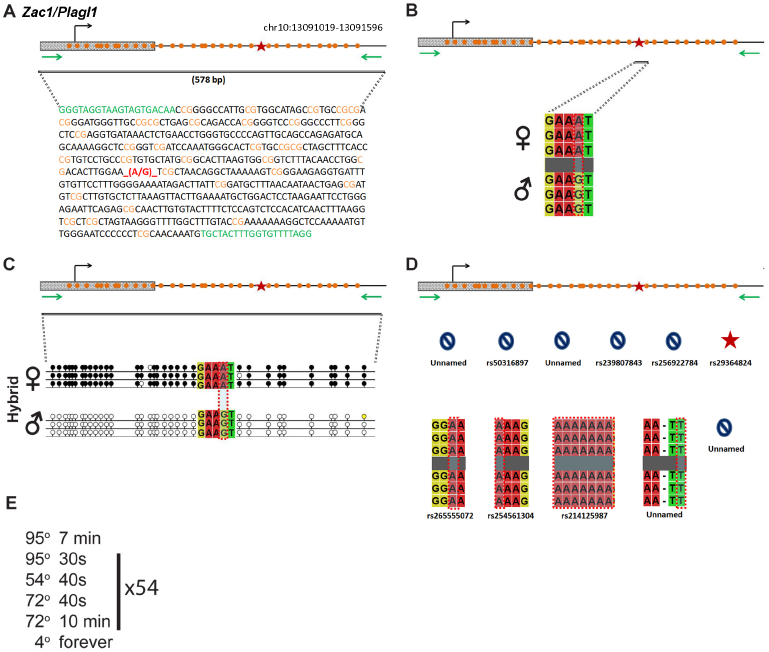
SNP verification within ZacliPlagil ICR. **A**. Schematic of Zaci/Plagli imprinting control region. Probed region is highlighted by double-dashed line with number of base pairs covered reported. CpG island indicated by dotted box. Green indicates primer sequences: orange indicates CpG dinucleotides: red star and bases indicate verified SNP. B. Verified SNP presented as sequences from B6 female and CAST male A-b-G SNP is highlighted by red dotted rectangle. **C**. Verification of proper imprinted status in hybrid 86/CAST progeny. SNP highlighted by red dotted rectangle. DNA methylation presented as lollipop diagram: white circles indicate unmethylated cytosines: black circles indicate methytated cytosines. **D**. Other SNPs reported in all three databases within the probed region with the SNP hlighted by red dotted rectangle. dbSNP identification number indicated under each SNP. Red star indicates validated SNP and blue closed circle indicates C-10-T polymorphism that cannot be assayed in bisulfite analysis. E. Optimal PCR conditions for probed region with the given primers.

### Summary of work

Of the SNPs that we analyzed we were able to validate 18, while we failed to validate 75 SNPs within those same regions (Table 1, red). In addition, a further 28 of them were C/T polymorphisms that bisulfite analysis was unable to differentiate (Table 1, blue). We also identified a SNP in the *Snrpn* ICR, which was not present in any of the three databases (Table 1, orange). Furthermore, during our optimization we failed to validate multiple SNPs that lie outside of our bisulfite primers. These SNPs are reported in Figure S1. Among the many SNPs reported in the dbSNP database that we failed to verify, most were identified as SNPs between strains other than CAST in the Sanger database. In the end, we could only find one SNP that was supposed to show a polymorphism based on the reported data but did not in our experiments (Table 1, purple). Thus, in general, we recommend using the Sanger database. However, it is important to note that since the Sanger database primarily contains SNPs located close or within genes, certain ICR SNPs had to be identified in the dbSNP database.

**Table 1:**
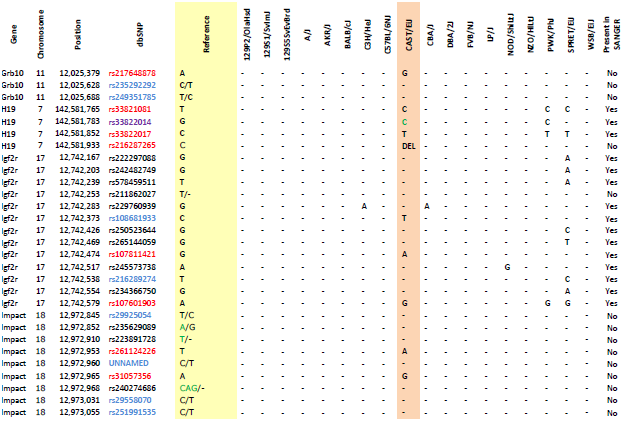
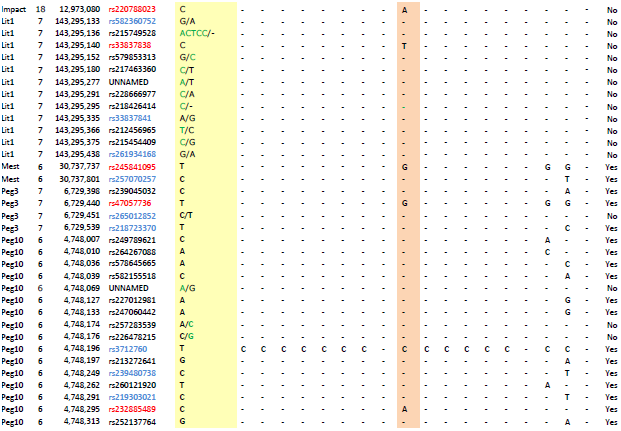
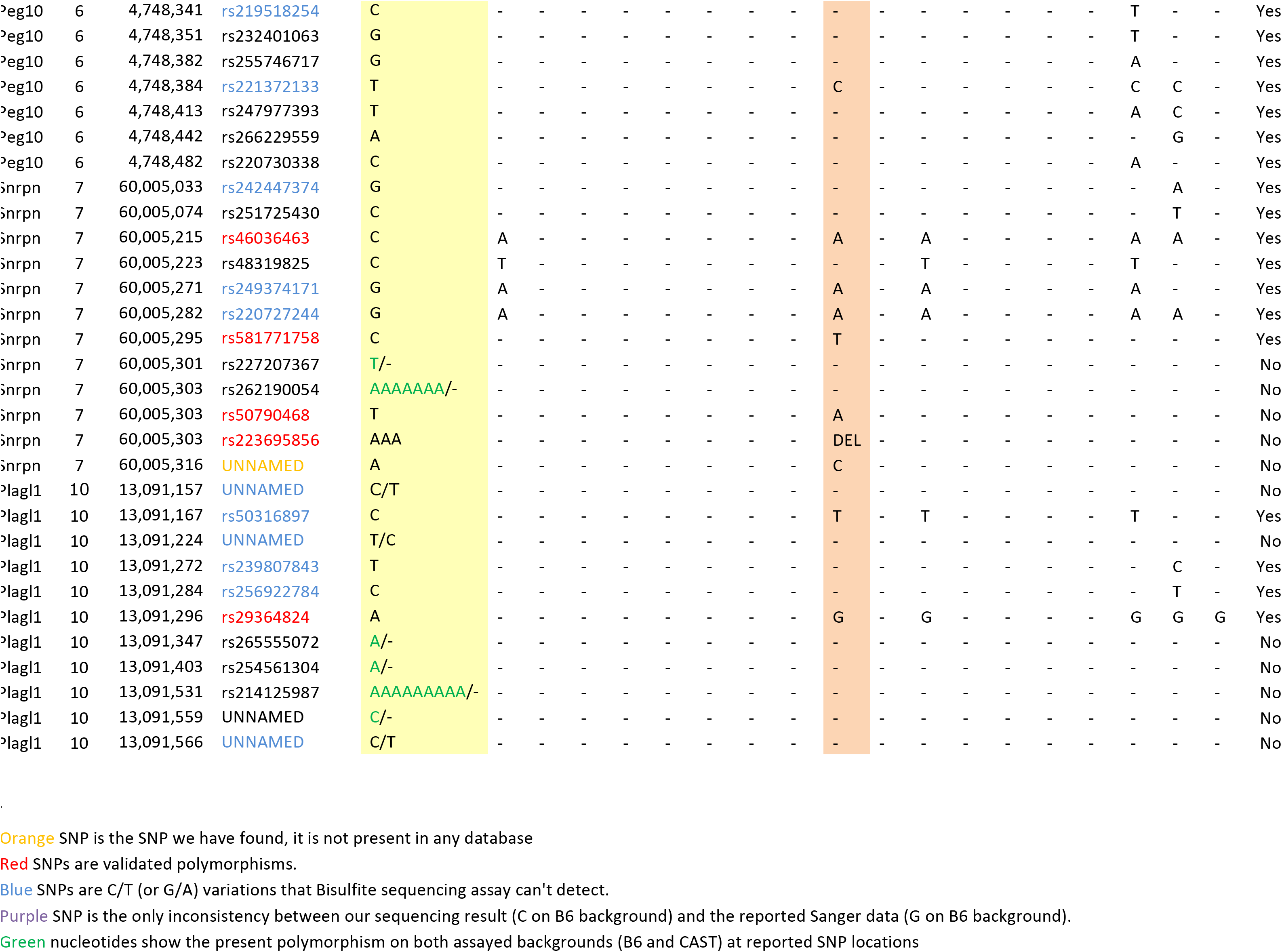
**The complete list of all the SNPs from 3 databases within surveyed regions**

In this resource, we have validated a number of SNPs within the ICRs of the most commonly imprinted loci. In addition, we have demonstrated a high frequency of invalid SNPs within ICRs when the pooled SNPs from the dbSNP (European variation archive) are used alone, highlighting the drawbacks of the mixed strain databases compared to the Sanger strain specific polymorphism database. Using the validated SNPs, we have optimized allele-specific DNA methylation assays that will allow for the rapid analysis of multiple imprinted loci in a variety of contexts, including at several ICRs that are not contained within the Sanger database. This resource will enable the systematic analysis of multiple imprinted genes in a number of potential applications.

### Potential applications

As this resource offers extensive and straight-forward assays to interrogate the most commonly studied imprinted loci, it can be utilized across a number of fields. There are two major instances where we envision the utility of this resource. First, cases where a regulatory mechanism directly interacts with multiple imprinted loci. Second, cases where a mechanism either indirectly regulates many imprinted loci, or affects multiple imprinted loci by generally disrupting the epigenetic landscape.

Recently, a number of proteins have been demonstrated to directly regulate multiple imprinted loci. These include, but are not limited to, *Dnmt3l, Dnmt1, Lsd2, Trim28, Zfp57*, and *Tet1/2*, each with a different mechanism of action (Bourc’his *et al.* 2001; Howell *et al.* 2001; Reik *et al.* 2003; Li *et al.* 2008a; Karytinos *et al.* 2009; Fang *et al.* 2010; Messerschmidt *et al.* 2012; Yamaguchi *et al.* 2013; Canovas and Ross 2016). For example, deletion of the regulatory subunit of the *de novo* DNA methyltransferase *Dnmt3L* results in the failure to establish maternal DNA methylation at a number of maternally imprinted loci, including *Peg3, Lit1/Kcnq1ot1* and *Snrpn* (Bourc’his *et al.* 2001; Hata *et al.* 2002). Another maternal effect enzyme required for the establishment of DNA methylation at maternally imprinted loci is the histone demethylase *Lsd2*. Mechanistically, *Lsd2* is required to remove H3K4 methylation in order to get proper DNA methylation at imprinted loci including *Mest, Grb10, and Zac1* (Ciccone *et al.* 2009; Lei *et al.* 2009; Fang *et al.* 2010; Zhang *et al.* 2012; Stewart *et al.* 2015). Furthermore, *Zfp57,* a KRAB domain zinc-finger protein, is required both maternally and zygotically to maintain the imprinting status of various imprinted loci including *Snrpn* (Li *et al.* 2008a; Strogantsev and Ferguson-smith 2012; Strogantsev *et al.* 2015). This protein is thought to bind directly to DNA with its zinc fingers and subsequently recruit factors that repress transcription (Li *et al.* 2008b; Quenneville *et al.* 2011; Strogantsev *et al.* 2015). These studies demonstrate how disruptions in mechanistically distinct regulatory mechanisms can affect multiple imprinted loci.

Alternatively, a number of mechanisms have been demonstrated to indirectly affect imprinted loci via general epigenetic disruptions. For example, mutations in human NLRP genes, which are required maternally for the transition to zygotic gene expression, result in hydatidiform moles and loss of imprinting (Docherty *et al.* 2015). Another maternal effect gene, *Lsd1,* the homolog of *Lsd2,* is also maternally required at fertilization for the maternal to zygotic transition (Ancelin *et al.* 2016; Wasson *et al.* 2016). Loss of maternal *Lsd1* leads to a general disruption of DNA methylation in the resulting progeny at both maternally and paternally imprinted loci (Ancelin *et al.* 2016; Wasson *et al.* 2016). These studies demonstrate how maternal factors, deposited into the zygote from the mother, are required for proper imprinting and development of the embryo.

As imprinting control regions are inherently asymmetric in their epigenetic modifications and opposing mechanisms are required at each parental ICR, even slight disturbances in the epigenetic landscape can lead to significant changes in expression at these loci. For example, disruptions in the maternal expression of *Grb10* results in developmental defects in mice, while disruption of the paternal allele of *Grb10* leads to changes in behavior, including increased social dominance (Garfield *et al.* 2011; Dent and Isles 2014). This highlights differences in the roles of imprinted parental alleles in mice. Another study that highlights the relative contributions of each parental allele describes parental specific duplications of the 15q11.2-q13.3 region of human chromosome 15 (Isles *et al.* 2016). Paternal duplications were more associated with autism spectrum disorder and developmental delay, while maternal duplications were more associated with psychiatric disorders (Isles *et al.* 2016). These studies demonstrate the complexity of outcomes associated with maternal versus paternal inheritance.

Finally, mechanisms that affect imprinted genes indirectly though general epigenetic disruptions highlight how the methylation status of ICRs can act as a proxy for global epigenetic alterations. For example, studies have demonstrated hypomethylation of a differentially methylated region in the *Igf2-H19* locus in Wilms tumor patients (Scharnhorst *et al.* 2001). In addition, embryos conceived using artificial reproductive technologies have higher incidences of Prader-Willi and Angelman Syndromes (Horsthemke and Wagstaff 2008; Buiting 2010; Butler 2011). These syndromes are caused by large-scale chromosomal abnormalities that affect multiple imprinted loci (Horsthemke and Wagstaff 2008; Buiting 2010; Butler 2011). It is also possible that imprinting may be disrupted by environmental factors. For example, Bisphenol A (BPA), an environmental toxin, as well as various endocrine disruptors, have been revealed to significantly alter the epigenetic landscape (Kang *et al.* 2011; Susiarjo *et al.* 2013). Also Vinclozolin exposure in mice leads to infertility due to sperm defects in mice which correlate with global alterations in the DNA methylation landscape (Anway *et al.* 2005; Kang *et al.* 2011). These studies demonstrate additional mechanisms that may lead to broad imprinting disruptions.

Due to various mechanisms that can disrupt the epigenetic landscape, we anticipate a growing need to assay imprinted loci in different mouse models. The resource provided here will facilitate the future analysis of multiple imprinted loci in a single hybrid genetic background.

## Acknowledgements

We would like to thank the epigenetic community at Emory University for their feedback. T. Lee for help in editing this manuscript and A. Ferguson-Smith and M. Bartolomei for feedback on the manuscript. In addition we would like to thank M. Bartolomei for providing the *H19* assay and D. Cutler for bioinformatics assistance. J.A.W was supported by the Biochemistry, Cell and Molecular Biology Training Grant (5T32GM008367). The work was supported by a grant to D.J.K from the National Science Foundation (IOS1354998).

